# Assessing sub-cellular resolution in spatial proteomics experiments

**DOI:** 10.1101/377630

**Authors:** Laurent Gatto, Lisa M. Breckels, Kathryn S. Lilley

## Abstract

The sub-cellular localisation of a protein is vital in defining its function, and a protein’s mis-localisation is known to lead to adverse effect. As a result, numerous experimental techniques and datasets have been published, with the aim of deciphering the localisation of proteins at various scales and resolutions, including high profile mass spectrometry-based efforts. Here, we present a meta-analysis assessing and comparing the sub-cellular resolution of 29 such mass spectrometry-based spatial proteomics experiments using a newly developed tool termed *QSep*. Our goal is to provide a simple quantitative report of how well spatial proteomics resolve the sub-cellular niches they describe to inform and guide developers and users of such methods.

## 1 Introduction

In biology, the localisation of a protein to its intended sub-cellular niche is a necessary condition for it to assume its biological function. Indeed, the localisation of a protein will determine its specific biochemical environment and its unique set of interaction partners. As a result, the same protein can assume different functions in different biological contexts. Protein mis-localisation can lead to adverse effects and have been implicated in multiple diseases [24, 5, 25].

Spatial proteomics is the systematic and high-throughput study of protein sub-cellular localisation. A wide range of techniques (reviewed in [10, 27]) and computational methods [11] have been documented, that confidently infer the localisation of thousands of proteins. Most techniques rely on some form of sub-cellular separation, many employing differential centrifugation or separation along density gradients, and the subsequent quantitative assessment of relative protein occupancy profiles in these sub-cellular fractions. Reciprocally, a broad array of computational methods have been applied, ranging from unsupervised learning e.g. clustering [29] and dimensionality reduction, and supervised learning such as classification (reviewed in [11]), semi-supervised learning and novelty detection [2] and, more recently, transfer learning [3] and Bayesian modelling [6].

Despite these advances, there has been a surprising lack of discussion in the community as to what constitutes a reliable spatial proteomics experiment, i.e a dataset that generates confident protein localisation results. It is however implicit that reliability and trust in the results is dependent on adequate sub-cellular resolution, i.e. *enough* separation between the different sub-cellular niches being studied to be able to confidently discern protein profiles originating from different sub-cellular niches. There are however currently no tools that enable estimating whether an experiment has been adequately resolved, and that would help data producers and consumers, to assess the adequacy of a dataset.

The importance of adequate sub-cellular resolution reaches beyond the generation of reliable static spatial maps. It is a necessary property of the data to consider tackling more subtle sub-cellular patterns such as multi-and trans-localisation, i.e. the localisation of proteins in multiple sub-cellular niches and the relocation of proteins upon perturbation [11].

In this work, we first describe how to understand and interpret widely used dimensionality reduction methods and visualise spatial proteomics data to critically assess their resolution. We then propose a simple, yet effective method, to quantitatively measure sub-cellular resolution and compare it across different experiments. We introduce QSep, a tool to assess the resolution of sub-cellular niches within spatial proteomics experiments, giving a metric of success of desired sub-cellular separations. Our approach fills an important gap in analysis of spatial proteomics data and will be useful to spatial proteomics practitioners to (1) assess the sub-cellular resolution of their experiments, (2) compare it to similar studies, (3) help set up and optimise their experiments, and additionally (4) to aid biologists in critically assessing spatial proteomics studies and their claims.

All the data and software used in this work are available in the `pRoloc` and `pRolocdata` packages [12]. The code to reproduce all results and figures presented here is available in the source of the reproducible document, available in the manuscript’s public repository [9] available at https://github.com/lgatto/QSep-manuscript/.

## 2 Spatial proteomics datasets

For this meta-analysis, we make use of 29 spatial proteomics datasets, summarised in table 1. These data represent a diverse range of species, sample types, instruments and quantification methodologies.

**Table 1:** Summary of the datasets used in this study. The percentage of variance along the principal components (PC) is related to the PCA plots on figure 14. All datasets are available in the `pRolocdata` package.

| Data | Proteins | Fractions | Clusters | PC var (%) | Title |
| --- | --- | --- | --- | --- | --- |
| hyperLOPIT2015 | 5032 | 20 | 14 | 72.26 | Protein and PMS-level hyperLOPIT datasets on Mouse E14TG2a embryonic stem cells from Christoforou et al. (2016). [4] |
| hyperLOPIT2015ms2 | 7114 | 10 | 14 | 74.72 | Protein and PMS-level hyperLOPIT datasets on Mouse E14TG2a embryonic stem cells from Christoforou et al. (2016). [4] |
| HEK293T2011 | 1371 | 8 | 12 | 65.04 | LOPIT experiment on Human Embryonic Kidney fibroblast HEK293T cells from Breckels et al. (2013) [2] |
| hirst2018 | 2046 | 15 | 12 | 82.50 | Data from Hirst et al. 2018 [15] |
| hyperLOPITU2OS2017 | 5020 | 40 | 12 | 63.29 | 2017 and 2018 hyperLOPIT on U2OS cells [28] (all fractions) |
| hyperLOPITU2OS2017b | 5020 | 37 | 12 | 67.68 | 2017 and 2018 hyperLOPIT on U2OS cells [28] (cleaned) |
| itzhak2016stcSILAC | 5265 | 30 | 12 | 70.61 | Data from Itzhak et al. (2016) [17] |
| itzhak2017 | 9201 | 30 | 12 | 31.39 | Data from Itzhak et al. 2017 [18] |
| rodriguez2012r1 | 2215 | 11 | 12 | 38.95 | Spatial proteomics of human inducible goblet-like LS174T cells from Rodriguez-Pineiro et al. (2012) [23] |
| tan2009r1 | 888 | 4 | 11 | 88.49 | LOPIT data from Tan et al. (2009) [26] |
| E14TG2aS1 | 1109 | 8 | 10 | 65.98 | LOPIT experiment on Mouse E14TG2a Embryonic Stem Cells from Breckels et al. (2016) [3] |
| trotter2010 | 347 | 16 | 10 | 81.14 | LOPIT data sets used in Trotter et al. (2010) [30] |
| beltran2016HCMV120 | 2045 | 6 | 9 | 79.36 | Data from Beltran et al. 2016 [19] HCMV infection, 120 hpi (hours post-infection) |
| beltran2016HCMV24 | 2196 | 6 | 9 | 77.18 | Data from Beltran et al. 2016 [19] HCMV infection, 24 hpi |
| beltran2016HCMV48 | 2206 | 6 | 9 | 74.95 | Data from Beltran et al. 2016 [19] HCMV infection, 48 hpi |
| beltran2016HCMV72 | 2062 | 6 | 9 | 75.32 | Data from Beltran et al. 2016 [19] HCMV infection, 72 hpi |
| beltran2016HCMV96 | 1868 | 6 | 9 | 76.59 | Data from Beltran et al. 2016 [19] HCMV infection, 96 hpi |
| beltran2016MOCK120 | 1757 | 6 | 9 | 71.84 | Data from Beltran et al. 2016 [19] MOCK, 120 hpi |
| beltran2016MOCK24 | 2220 | 6 | 9 | 80.12 | Data from Beltran et al. 2016 [19] MOCK, 24 hpi |
| beltran2016MOCK48 | 2181 | 6 | 9 | 68.90 | Data from Beltran et al. 2016 [19] MOCK, 48 hpi |
| beltran2016MOCK72 | 2161 | 6 | 9 | 68.83 | Data from Beltran et al. 2016 [19] MOCK, 72 hpi |
| beltran2016MOCK96 | 1748 | 6 | 9 | 73.24 | Data from Beltran et al. 2016 [19] MOCK, 96 hpi |
| dunkley2006 | 689 | 16 | 9 | 86.70 | LOPIT data from Dunkley et al. (2006) [7] |
| foster2006 | 1555 | 26 | 8 | 53.13 | PCP data from Foster et al. (2006) [8] |
| nikolovski2014 | 1385 | 20 | 8 | 67.97 | LOPIMS data from Nikolovski et al. (2014) [22] |
| groen2014cmb | 424 | 18 | 7 | 64.30 | LOPIT experiments on Arabidopsis thaliana roots, from Groen et al. (2014) [13] |
| nikolovski2012imp | 1385 | 32 | 7 | 77.82 | Meta-analysis from Nikolovski et al. (2012) [21] |
| andreyev2010rest | 2642 | 36 | 6 | 25.39 | Six sub-cellular fraction data from mouse macrophage-like RAW264.7 cells from Andreyev et al. (2009) [1] |
| hall2009 | 1090 | 16 | 5 | 63.45 | LOPIT data from Hall et al. (2009) [14] |

We have applied minimal post-processing to the data and have used, as far as possible, the data and annotation provided by the original authors. The data from Foster et al. [8] have been annotated using the curated marker list from Christoforou et al. [4], as only a limited number of markers were provided by the authors^1^. Marker proteins are well-known and trusted, generally manually curated residents of sub-cellular niches, for the species and condition of interest, and are used to annotate the spatial proteomics experiment. These annotations are then used for visualisation and quality control (see section 3.2) and supervised machine learning.

We have also only considered sub-cellular niches (also referred to as protein clusters, or clusters) that were defined by at least 7 protein markers^2^. Where studies include multiple replicated experiments, we used the combined dataset, rather than using the individual replicates, as combining data often leads to better sub-cellular resolution [30]. In addition, for dimensionality reduction and visualisation, we have systematically replaced missing values by zeros. When calculating distances between protein profiles (see section 3.3), however, missing values were retained.

It is important to highlight that not all experiments used in this study have as a main goal the generation of a global (or near global) sub-cellular map. While the works Christoforou et al. [4] (*A draft map of the mouse pluripotent stem cell spatial proteome*) and Itzhak et al. [17] (*Global, quantitative and dynamic mapping of protein subcellular localization*) explicitly state such goal, other experiments such as Groen et al. [13] (*Identification of trans-golgi network proteins in Arabidopsis thaliana root tissue*) or Nikolovski et al. [22] (*Label free protein quantification for plant Golgi protein localisation and abundance*) have a more targeted goal (identification of trans-Golgi-network and Golgi apparatus proteins, respectively). When an experiment is targeted at resolving one or two particular niche, it is often the case that other sub-compartments are less well-resolved and hence, it is important to keep the overall aim of the studies in mind when assessing their overall resolution.

## 3 Assessment

While never performed in a systematic way as in the work presented here, some authors have provided some metrics to demonstrate the quality of their data. One such metric is the macro-F1 score, calculated during classification parameter optimisation (see the `pRoloc` [12] documentation or [11] for details about this procedure). Briefly, the macro-F1 is computed as the harmonic mean of the precision (a measure of exactness) and recall (a measure of completeness) on marker proteins to infer a set of good model parameters to be used subsequently, when inferring the localisation of proteins of unknown location. The scores are computed during a number of iterations, where some markers are used as part of a validation partition, and others are used for parameter selection using cross-validation. These optimisation metrics are computed over a range of parameters on different subsets of marker proteins, and optimal parameters are optimised for the subset of markers used at each iteration, and hence are likely to provide an over-fitted view of the data. Note that while each iteration is over-fitted, the goal of the iterative procedure is to identify recurring best parameters, in the hope that these will subsequently generalise to new, non-marker proteins.

One study [17] used the Pearson correlation between replicates to demonstrate the replicability of their experiments. While useful within an experiment, correlation cannot be compared between experiments, as the correlation values will depend on the number of proteins in the experiment. More generally, correlation isn’t a good measure of reproducibility [16]; it focuses on values without context. A better here approach would be to compare classification results and demonstrate that these agree across different replicates.

### 3.1 Sub-cellular diversity

A first assessment that provides an important indication of the resolution of the data concerns the number and diversity of sub-cellular niches that are annotated. In the 29 datasets used in this study, this number ranged from 5 (dataset *hall2009*) to 14 (dataset *hyperLOPIT2015*). These numbers should be assessed in the light of about 25 different sub-cellular niches that are documented in all 29 datasets, which are still underestimating the biological sub-cellular complexity. The Human Protein Atlas [28] (https://www.proteinatlas.org), for example, classified 33 sub-cellular organelles and fine structures.

### 3.2 Dimensionality reduction and visualisation

Principal component analysis (PCA) is a widely used dimensionality reduction technique in spatial proteomics. It projects the protein occupancy profiles into a new space in such a way as to maximise the spread of all points (i.e. marker proteins and proteins of unknown localisation) along the first new dimension (principal component, PC). The second PC is then chosen to be orthogonal to the first one while still maximising the remaining variability, and so on. Each PC accounts for a percentage of the total variability and it is not uncommon, in well executed experiments, that the two first PCs summarise over 70% of the total variance in the data, confirming that the resulting visualisation remains a reliable and useful simplification of the original, multidimensional data.

By firstly summarising the occupancy profiles along PC1 and PC2 (and, possibly, other PCs of interest if necessary), it becomes possible to visualise the complete dataset in a single figure (as opposed to individual sets of profiles - see for example figure 5 in Gatto et al. [10]). In a first instance, it is advised to visualise the data without marker annotation to confirm the presence of discrete clusters, i.e. dense clouds of points that are well separated from the rest of the data (see for example data from Christoforou et al. [4] on figure 1, left). Such patterns can further be emphasised by using transparency (figure 1, centre) or binned hexagon plots (figure 1, right) to highlight density.

**Figure 1:**
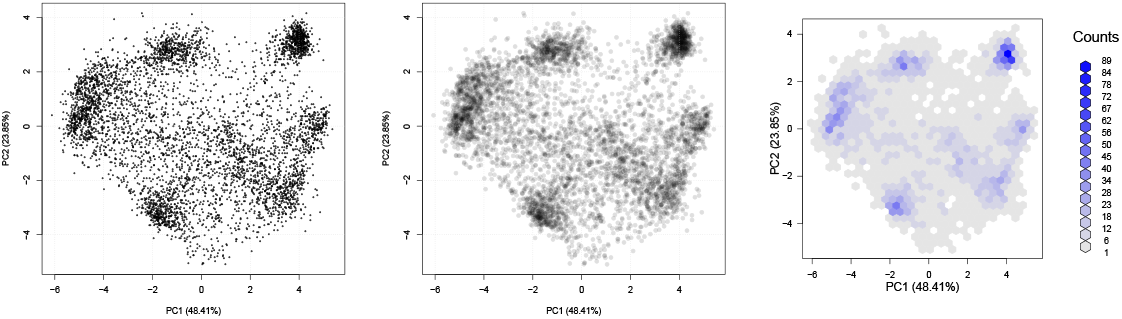
Unsupervised visualisation of spatial resolution using the `plot2D` function from the `pRoloc` package.

In figure 2, three datasets are compared to illustrate different levels of cluster density and separation. We see areas of high density (many proteins per hexagon), that are highlighted by dark blue bins, and less dense areas, displayed as white/grey regions, as indicated by the count key on the right-hand side of each plot. The figure on the left is the hyperLOPIT data from Christoforou et al. [4] (as on figure 1) that used synchronous precursor selection (SPS) MS^3^ on an Orbitrap Fusion. The middle figure represents the same experiment and same proteins, analysed using conventional MS^2^, illustrating the effect of reduced quantitation accuracy. Finally, on the right, an experiment with considerable less resolution [14].

**Figure 2:**
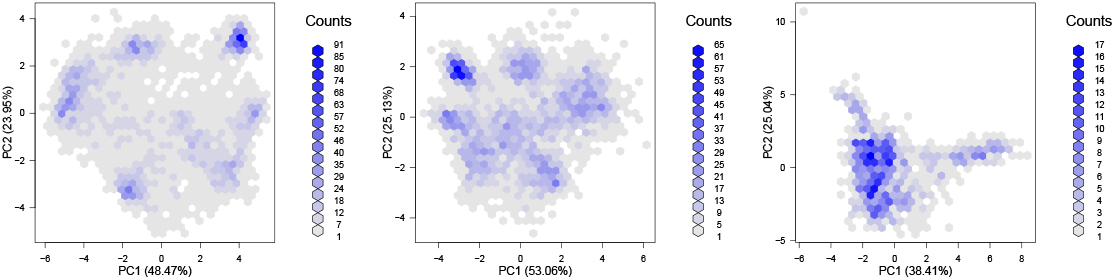
Comparing the cluster density and separation of experiments with excellent (left), intermediate (centre) and poor (right) resolution.

Considering that the aim of sub-cellular fractionation is to maximise separation of sub-cellular niches, one would expect sub-cellular clusters to be separated optimally in a successful spatial proteomics experiment. In PCA space, this would equate to the location of the marker clusters along the periphery of the data points. In other words, the maximum variability of a successful spatial proteomics experiments should be reflected by the separation of genuine (i.e. expected/annotated marker) spatial clusters, as illustrated on figure 3.

**Figure 3:**
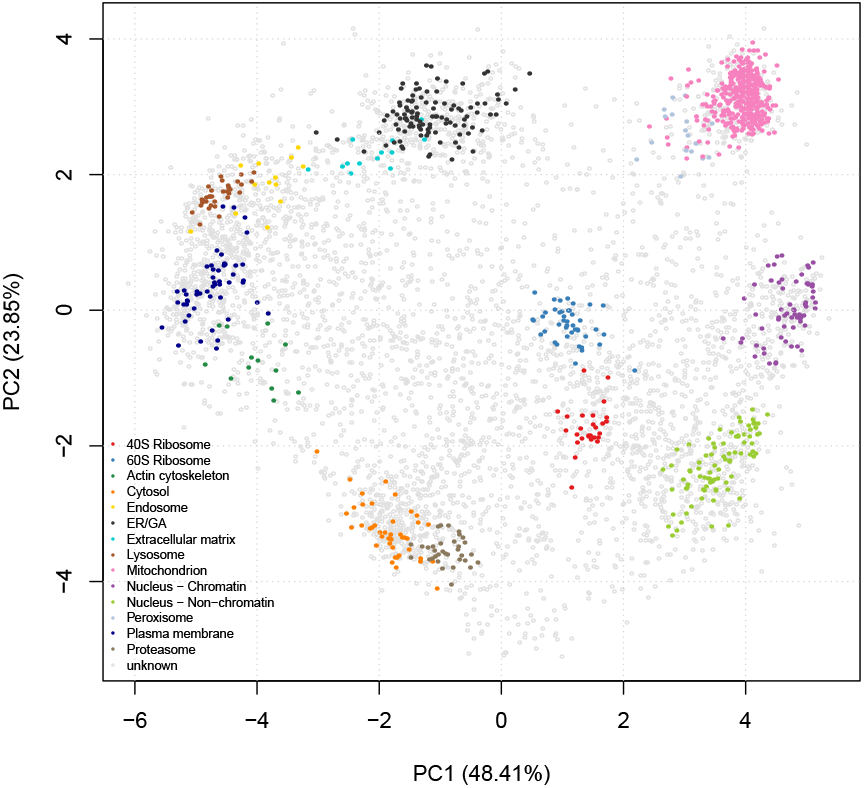
Annotated PCA plot of the *hyperLOPIT2015* dataset.

Another dimensionality reduction method of interest is linear discriminant analysis (LDA). LDA will project the protein occupancy profiles in a new set of dimensions using as a criterion the separation of marker classes by maximising the between class variance to the within class variance ratio. As opposed to the *unsupervised* PCA, the *supervised* LDA should not be used as an experiment quality control, but can be useful to assess if one or more organelles have been preferentially separated. LDA and many other dimensionality reduction techniques (such as t-SNE [31] for instance) are readily available in `pRoloc`.

It is important to highlight that these representations, while generally reflecting a major proportion of the variability in the data, are only a summary of the total variability. Some sub-cellular niches that overlap in 2 dimensions can be separated along further components. It is sometimes useful to visualise data in three dimensions (using for example the `plot3D` function in the `pRoloc` package), which still, however, only reflect part of the total variability.

When assessing the resolution of some specific organelles of interest, one should compare the full protein profiles of the marker proteins as illustrated on figure 4 (`pRoloc`’s `plotDist` function can be used for this, or the interactive application `pRolocVis` in the `pRolocGUI` package), or visualise a dendrogram representing the average distance between cluster profiles (the `mrkHClust` function from `pRoloc` offers this functionality). While detailed exploration of a dataset using these and other visualisation approaches is crucial before analysing and interpreting a new spatial proteomics experiment, a detailed exploration of each of the 29 datasets used in this meta-analysis is out of the scope of the study presented here.

**Figure 4:**
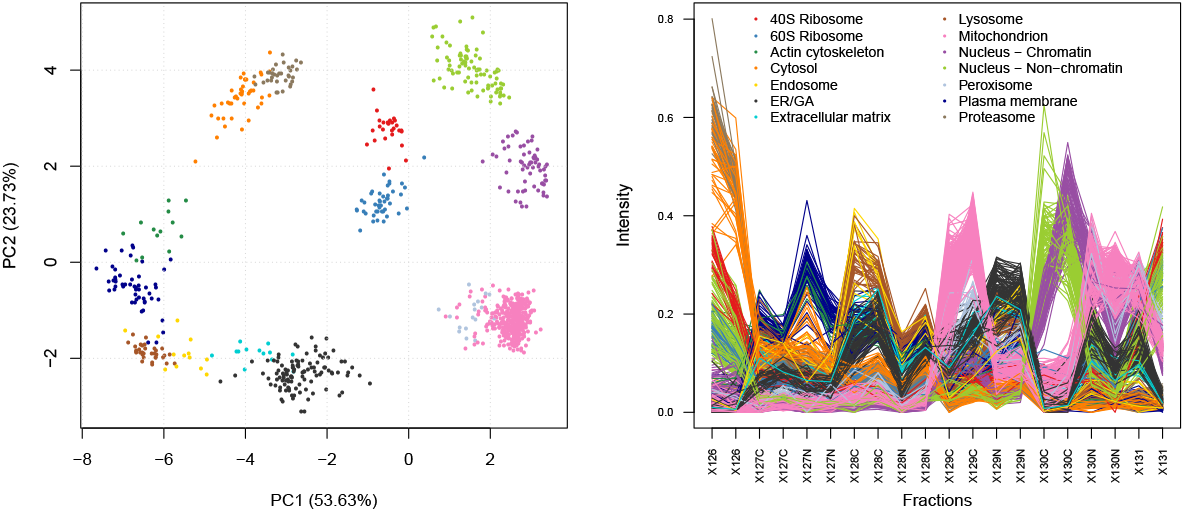
Marker proteins of the *hyperLOPIT2015* data set along PC 1 and 2 (left), and visualising the full proteins profiles (right). We see that while the mitochondrion and peroxisome clusters aren’t separated along the two first principal components (they are along PC 7), line plots displaying the n-dimensional protein profiles show that while similar, these two compartments differ along fractions X129C.

### 3.3 Quantifying resolution

While visualisation of spatial proteomics data remains essential to assess the resolution, and hence the success, of a spatial proteomics experiment, it is useful to be able to objectively quantify the resolution and directly compare different experiments. Here, we present a new infrastructure, termed QSep, available in the `pRoloc` package [12], to quantify the separation of clusters in spatial proteomics experiments. It relies on the comparison of the average Euclidean distance of the full, n-dimensional protein profiles *within* and *between* sub-cellular marker clusters. As illustrated on the heatmaps in figure 5 for the *hyperLOPIT2015* data, these distances always refer to one reference marker cluster.

**Figure 5:**
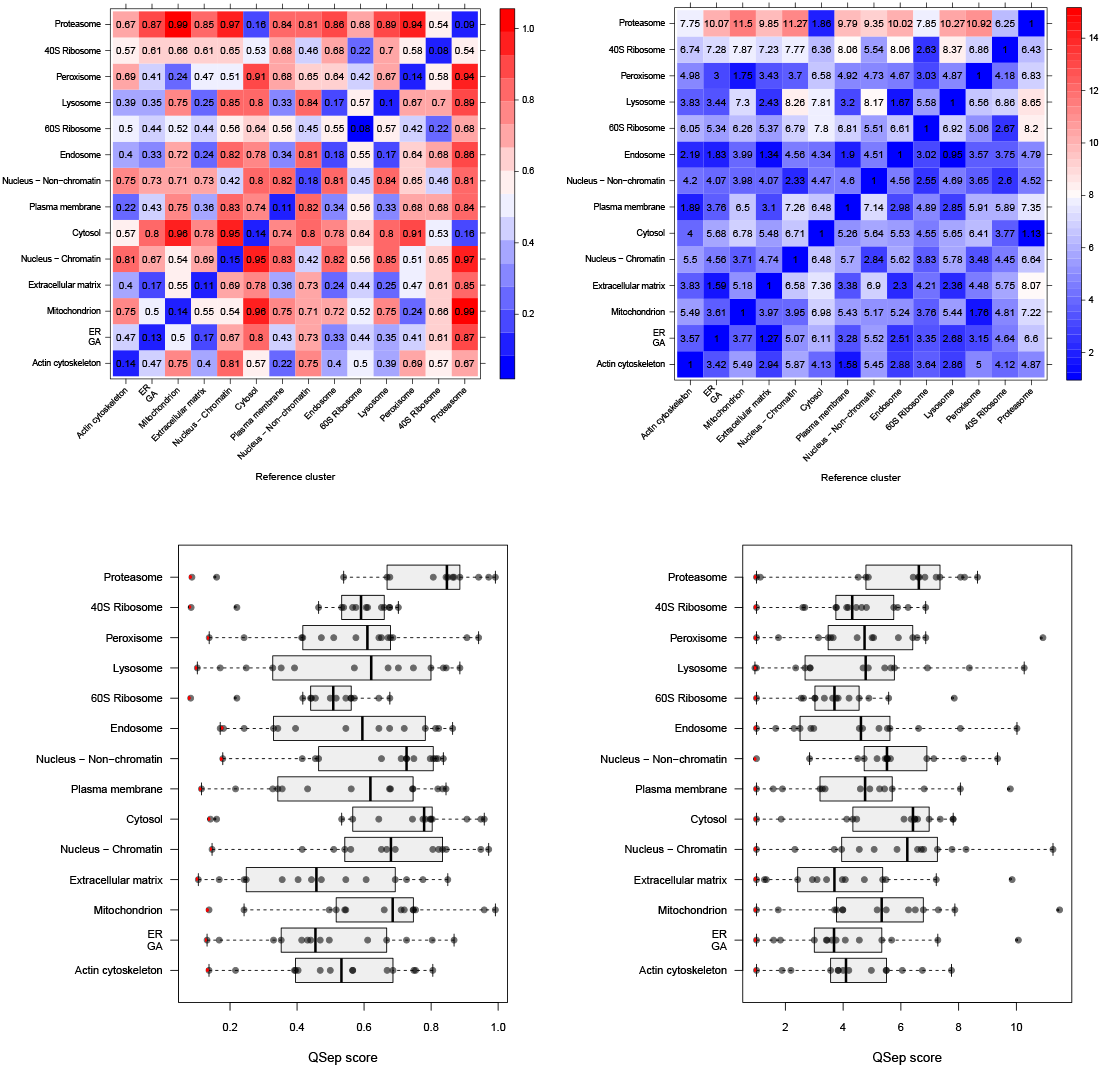
Quantifying resolution of the *hyperLOPIT2015* data Christoforou et al. [4]. The heatmaps at the top illustrate the raw (left) and average normalised (right) within (along the diagonal) and between euclidean cluster distances. The boxplots at the bottom summarise these same values (raw on the left, normalised on the right) to enable easier comparison between clusters, where the within distances are highlighted in red.

The raw distance matrix (figure 5, top-left) is symmetrical (i.e. the distance between cluster 1 and 2 is the same as between cluster 2 and 1). Within cluster distances, along the diagonal, are generally the smallest ones, except when two clusters overlap, as the lysosome and endosome in our example. To enable the comparison of these distances within and between experiments (see section 4 for the latter), we further divide each distance by the reference within cluster average distance (figure 5, top-right). This thus gives information about how much the average distance between cluster 1 and 2 is greater than the average distance within cluster 1 (i.e. the tightness of that cluster). At this stage, the distance matrix is no longer symmetrical. To facilitate the comparison of distances between organelles, the distance distributions can also be visualised as boxplots (figure 5, bottom).

The rational behind these measures is as follows. Intuitively, we assess resolution by contrasting the separation between clusters (formalised by the average distance between two clusters) and the tightness of single clusters (formalised by the average within cluster distance). Ideal sub-cellular fractionation would yield tight and distant clusters, represented by a large normalised between cluster distances on figure 5.

### 3.4 Application of the assessment criteria

To further demonstrate the interpretation of these resolution metrics, we directly compare the two recent global cell maps from [4] (dataset *hyperLOPIT2015*) and [17] (dataset *itzhak2016stcSILAC*). Both feature high protein coverage (7114 and 5265 proteins respectively) and good sub-cellular diversity (14 and 12 annotated clusters respectively). The former contains duplicated experiments, each made of 10 fractions and the latter contains 6 replicates with 5 fractions each. The *itzhak2016stcSILAC* dataset has been produced using the *Dynamic Organellar Maps* design, which goes some way to also determine relative abundances across proteins, an important feature that is however not considered here, when computing the QSep scores.

Figure 6 shows the PCA plots applying transparency to identify the underlying structure in the quantitative data and the annotated versions using the markers provided by the respective authors.

**Figure 6:**
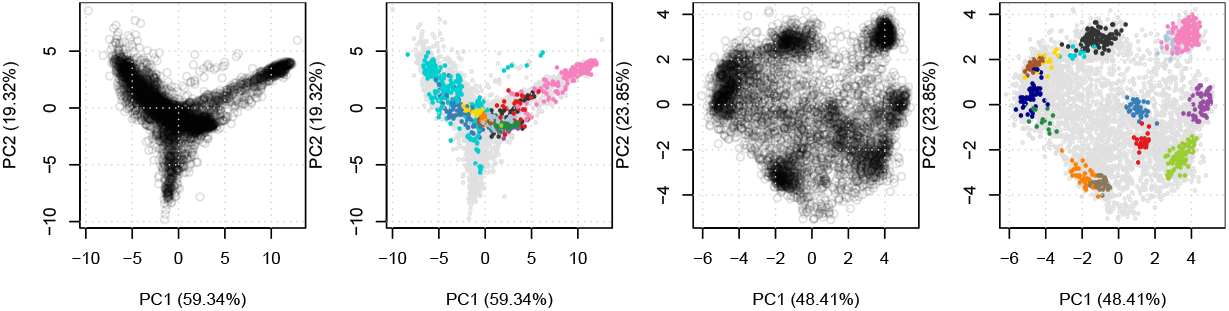
PCA plots for *itzhak2016stcSILAC* (left) and *hyperLOPIT2015* (right). Here, we display PC 1 and 2 for both datasets for comparability. The original authors displayed PC 1 and 3 for the *itzhak2016stcSILAC* data (see figures 13 and 14 below).

Figure 7 illustrates the normalised distance heatmaps and boxplots for the two datasets *(itzhak2016stcSILAC* at the top and *hyperLOPIT2015* at the bottom). The two heatmaps, that have been rendered using the same colour-scale, display strikingly different colour patterns. The top heatmap shows a majority of small normalised distances (dark blue cells) and with only a limited number of large distances (red cells), along the mitochondrial reference cluster. Conversely, the bottom heatmap displays a majority of average (light blue and white cells) and larger distances (red cells) across all sub-cellular clusters. The boxplots allow a more direct comparison of the distances across the two datasets. On the top boxplot, we detect relatively short distances for most clusters, with most large distances stemming from the mitochondria, leading to a median distance of 2.48. The distributions on the bottom boxplot show larger distances, equally spread among all clusters, with an median distance of 4.91.

**Figure 7:**
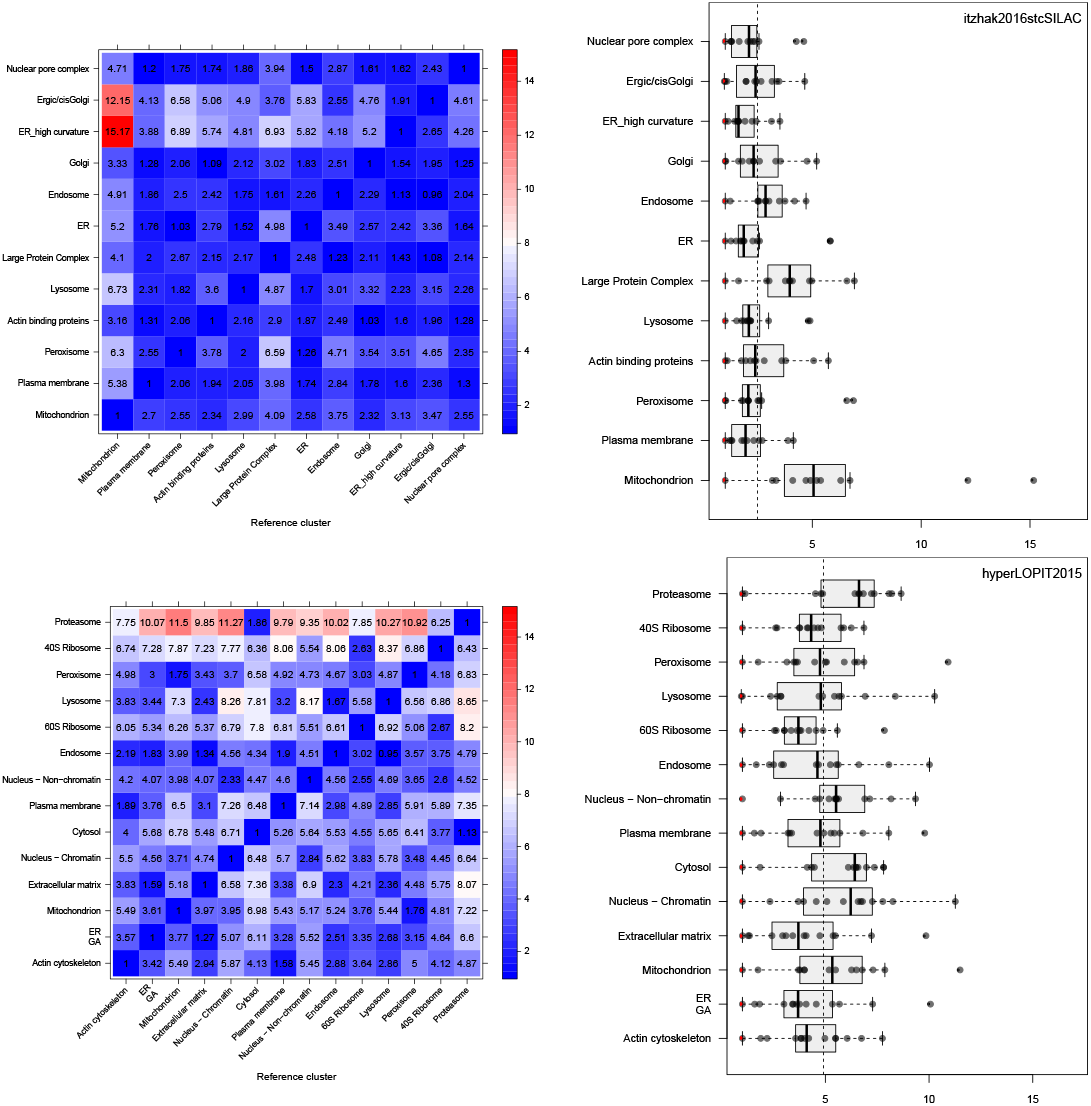
Contrasting quantitative separation assessment between the *itzhak2016stcSILAC* [17] (top) and *hyperLOPIT2015* [4] (bottom) datasets. The dashed vertical lines on the boxplots represent the overall median between cluster distance, 2.48 and 4.91 for *itzhak2016stcSILAC* and *hyperLOPIT2015* respectively.

## 4 Comparative study

We next apply the quantitative assessment of spatial resolution described in section 3.3 to compare the 29 experiments presented in section 2. Figure 8 shows, for each dataset, a boxplot illustrating the distribution of the global average normalised distances for all spatial clusters. The datasets have been ordered using the experiment-wide median between distance. It is important to always refer back to the original data when considering summarising metrics like these, to put the resolution into context. In particular, while high global resolution is essential for experiments that aim for a cell wide exploration, it will not be a useful metric in experiments that focus on a limited sub-cellular niches. The density and annotated PCA plots discussed in section 3.2 are provided in figures 13 and 14 and the quantitative assessment boxplots and heatmaps are shown in figures 15 and 16.

**Figure 8:**
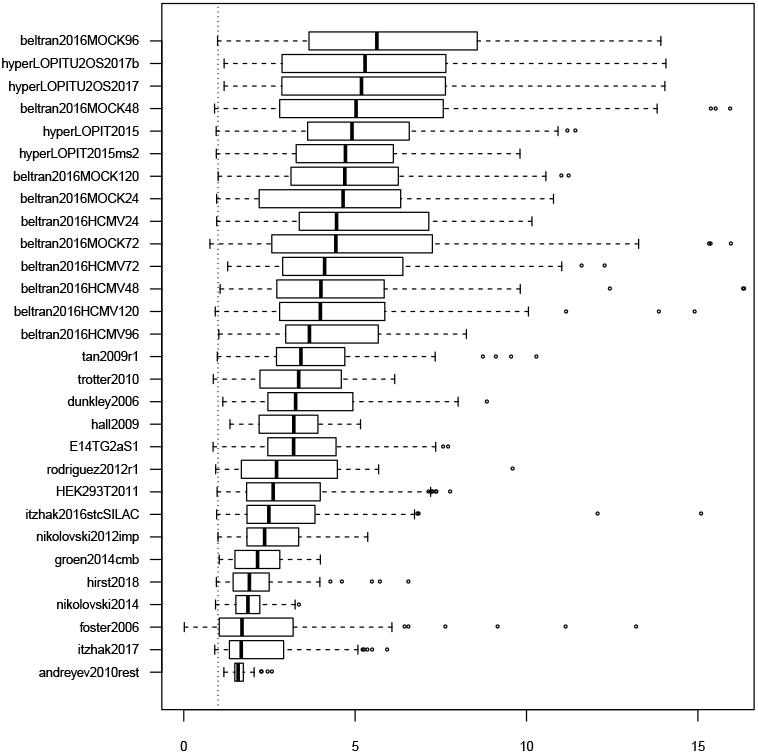
Quantitative separation assessment using experiment-wide normalised distances between cluster distances. The vertical line represents the normalised intra-cluster distance of 1.

The individual LOPIT-based experiments from Jean Beltran et al. [19] and the embryonic mouse stem cell [4] and U2OS [28] hyperLOPIT experiments (using SPS MS^3^ and conventional MS^2^) show the best experiment-wide resolution. The next set of experiment are *tan2009r1, trotter2010, dunkley2006, hall2009* and *E14TG2aS1*. It is important to highlight that most of the datasets (as well as *HEK293T2011*, discussed later) have either been directly re-analysed using a semi-supervised novelty detection algorithm *phenoDisco* [2] (the only exception here being *hall2009*), or, in the case of *trotter2010*, have been annotated using markers based on the *phenoDisco* re-analysis. The novelty detection algorithm, *phenoDisco*, searches for new clusters of unlabelled proteins, using the marker proteins to guide the clustering of unlabelled features. These new clusters, termed *phenotypes*, are then validated by the user for coherence with known sub-cellular niches. This re-analysis has proven successful [2] and has identified previously undetected sub-cellular niches that form tight and well-resolved clusters (see for example ribosomial and trans-Golgi network (TGN) in *dunkley2006*, or proteasome and nucleus in *tan2009r1* to cite only a few), which in turn favour good resolution scores. The *hall2009* dataset is relatively poorly annotated (only 5 sub-cellular clusters, which is the lowest in all test datasets). As long as these few clusters are well separated, poor annotation will however not negatively influence the resolution scoring.

The next set of experiments that show comparable resolution profiles are *HEK293T2011, itzhak2016stcSILAC* and *nikolovski2012imp*. Note that the quantitative separation measurement is robust to questionable marker annotation. For example, the *Lange Protein Complex* class defined by the original authors in the *itzhak2016stcSILAC* data could be dropped as it loosely defines many niches and thus lacks resolution. This omission only marginally influences the overall assessment metrics as only the distances to/from that class are affected (i.e. 23 out of 144 distances) and as such it would not change its rank among the test datasets.

As mentioned earlier, the *groen2014cmb* and *nikolovski2014* datasets are targeted experiments, focusing on the trans-Golgi-network and Golgi niches respectively. Such experiments do not aim for the best global resolution, which is reflected by relatively low global resolution.

The *foster2006* experiment displays relatively poor separation. This might be due to the relatively high number of missing values (42.4 %). Finally, the *andreyev2010rest* dataset suffers from very broad sub-cellular clusters (compared to separation between clusters).

The PCA plots and QSep heatmaps for all datasets are provided in the appendix, section A.

## 5 Assessing the resolution metric

Next, we assess the resolution metric, and how the annotation of the spatial proteomics data influences the metric itself.

We find that the number of classes does not have any effect on the resolution assessment scoring. Indeed, dropping any class will result in a sub-sample of normalised inter-cluster distances, with random variations around the overall median inter-cluster distance. In figure 9, we show the distribution of the resolution metrics when removing all possible combinations of 1 to 3 sub-cellular classes for the *E14TG2aS1* dataset, that displays an average overall resolution, and *hyperLOPIT2015*, that has among the highest resolution. In both cases, we see that the number of removed classes does not influence the overall score distributions. When modelling the linear relation between the median scores and the number of removed classes, the slopes are 0.036 and 0.01 respectively.

**Figure 9:**
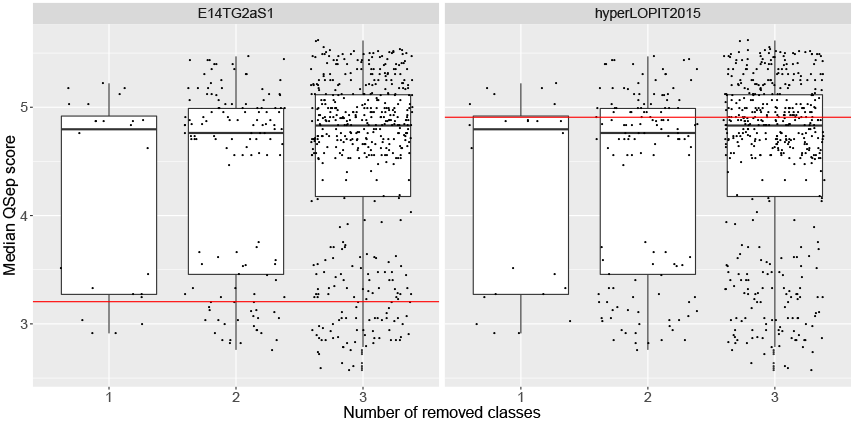
Effect of removing sub-cellular clusters on the resolution metric for the *E14TG2aD1* (left) and *hyperLOPIT2015* experiments (right). Each dot represents a median resolution score for the experimental setting (i.e missing n classes). The horizontal lines represents the median resolution metrics for the complete dataset. Note the overall higher median assessment scores for the better *hyperLOPIT2015* experiment

The definition of marker proteins has of course an effect on the assessment metric. Tighter clusters will result in smaller intra-class distances and, as a result, in larger normalised inter-class distances. To illustrate the effect of marker definition, we transferred the marker annotation between the *hyperLOPIT2015* and *itzhak2016stcSILAC* datasets (see PCA plots on figure 10, left) and calculated the quantitative resolution metrics (see boxplots on figure 10, right).

**Figure 10:**
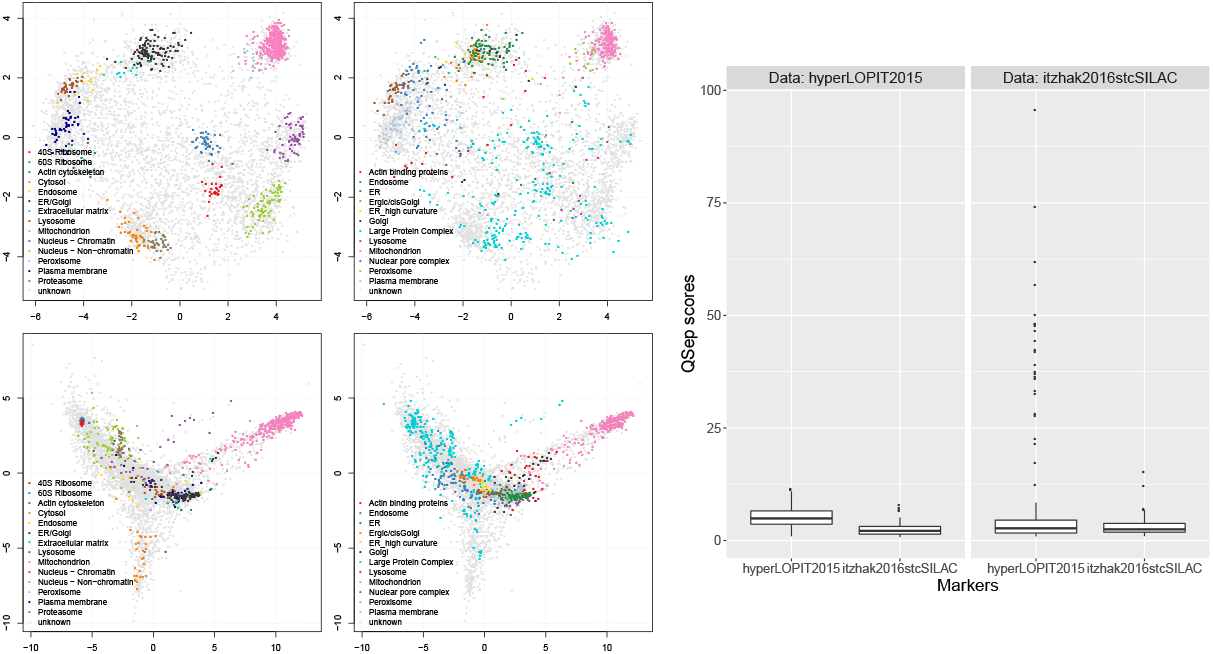
Marker transfer between *hyperLOPIT2015* and *itzhak2016stcSILAC*. On the left, the 4 data/marker combinations are displayed on PCA plots: the top and bottom row contains the *hyperLOPIT2015* and *itzhak2016stcSILAC* data respectively, while the left and right columns display the markers from *hyperLOPIT2015* and *itzhak2016stcSILAC* respectively. On the right, the resolution scores have been calculated for the same markers (along the *x* axis) data (left and right panels). The respective median scores are, from left to right, 4.91, 2.15, 2.71 and 2.48.

As expected from the PCA plots on figure 10 and the diffuse *Lange Protein Complex* cluster, and as can be seen on QSep distribution boxplots, transferring the *itzhak2016stcSILAC* markers to the *hyperLOPIT2015* dataset (distributions in the left panel) has a detrimental effect on the separation (testing the log-transformed distributions with a t-test produces a p-value of 4.3 × 10^−32^). Annotating *itzhak2016stcSILAC* with the *hyperLOPIT2015* markers (distributions in the right panel) hardly improves its resolution metric (p-value of 0.0015, log-transformed QSep scores). The main effect here is to emphasise the separation between the mitochondrion and other spatial niches, in particular the very tight 40S and 60S ribosomal clusters. These examples illustrate the importance and impact of marker curation and annotation for individual experiments. In particular, the *Large Protein Complex* cluster from *itzhak2016stcSILAC*, while also diffuse in its original dataset, has a severe effect on a dataset that it was not curated for.

## 6 Conclusions

In this manuscript, we have described in detail how to assess and quantify the resolution of spatial proteomics experiments. We have applied dimensionality reduction and visualisation, as well as a simple and intuitive quantitative metric to explore and compare a variety of publicly available spatial proteomics datasets using the annotation provided by the original authors. We have also assessed the resolution metric itself and observed that it was immune to the number of clusters used for its computation and showed the possible influence of different marker annotation on the metric itself.

The ordering of the quantitative resolution detailed in section 4 should not be taken as absolute. Its main purpose is to provide a guide to compare different experiments. It will be useful for laboratories that do spatial studies on different models and with different fractionation and/or quantitation methods, to assess the impact of these variables. It is also useful to compare resolution between different labs, as demonstrated in our comparative study (section 4). We anticipate that it will also prove useful for the researcher wanting to assess the resolution of newly published studies, and put them into a wider context. It is necessary to emphasise the importance and effect of marker definition and curation on estimating and assessing the resolution of spatial proteomics experiments (section 5) and, of course, the impact of markers on the subsequent assignment of proteins to their most likely sub-cellular compartments. Sub-cellular resolution is of course only one aspect of spatial proteomics, albeit an important one, that critically determines the reliability of protein assignments to spatial niches as well as the identification of multi- and trans-localisation events.

Finally, we reflect on the implications of this work on the spatial proteomics community, and more generally the cell biology community that relies on protein localisation data. We have assessed dataset spanning 12 years of spatial proteomics. Since 2006, the community has seen many important improvements: significant advances in mass spectrometry quantification methodologies (see figure 2 comparing SPS MS^3^ and conventional MS^2^ for an example), improvements in biochemical fractionation methods and spatial proteomics designs (as exemplified by the substantial improvement obtained by hyperLOPIT [4]), careful data annotation and marker curation, as well as considerable breakthroughs in data analysis(for example using semi-supervised learning [2] or a Bayesian spatial proteomics framework [6]). One might then wonder to what extend benefits have lead to tangible improvements in resolution over time?

On figure 11, we have ordered the datasets’ resolution metric according to their publication year. We can see that a set of recent datasets, including the mouse stem cell [4] and U2OS [28] hyperLOPIT experiments (published in 2016 and 2017 respectively), and [19] (published in 2016), a variation of the LOPIT experiment, show a superior resolution.

**Figure 11:**
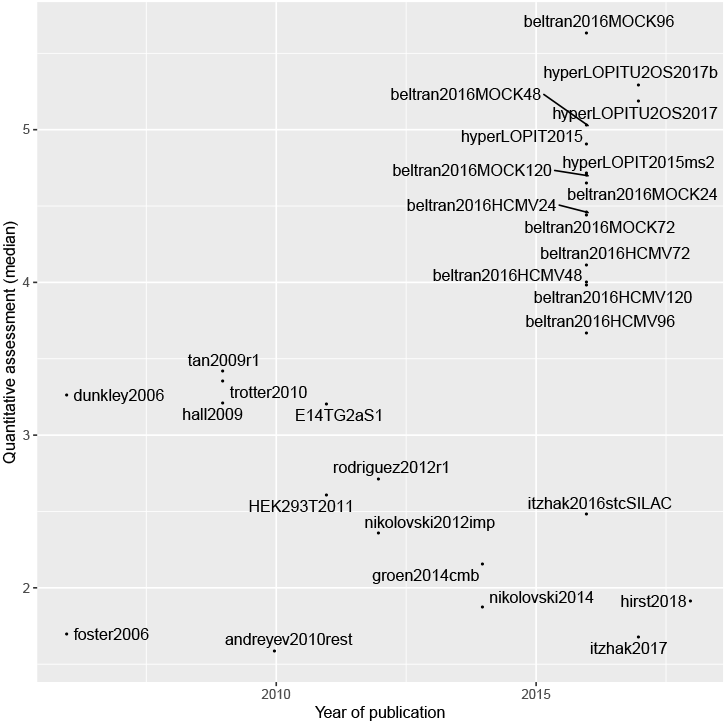
Resolution of spatial proteomics experiments over time. The assigned dates match either the original publication or, when know, the actual date the data was generated. For publications that re-analysed data, the date of the original publication of generation was used. When publications used several datasets, the date of the most recent one was used.

While the definition of sub-cellular resolution, as defined by the QSep measure, is only one aspect of spatial proteomics, one could argue that the community at large would benefit from a more systematic approach when considering the resolution of spatial proteomics experiments. It is however difficult to draw general conclusions whether any experimental properties correlate with better resolution due to the limited number of datasets (29 dataset, out of 18 individual publications, some including data re-analyses), as well as confounding factors such as the laboratory of origin and time of publication. As an example of the latter (figure 12, left panels), the distinction between iTRAQ and TMT tagging would falsely suggest that the latter (used in the high-performing `beltran` and `hyperLOPIT` experiments) is better, when this trend is purely the result of the field moving towards isobaric systems with more tags to quantify more fractions along the gradient (leading to better resolution [11]) irrespective of the quantitation technology. Similarly, no pattern based on the species is visible (figure 12, right panels). In our experience, sample-specific factors such as the heterogeneity of cells or tissue (such as for example the U2OS or HeLa cell lines used in [28] or [17] vs. *Drosophila* embryos in [26] or *Arabidopsis* root tissue in [13]), the efficiency of cell harvesting (in suspension vs. adherent cells) and membrane fractionation (requiring careful sample-specific optimisations) will have a crucial effect of the final resolution of the data. The species will however play an important role in the availability of reliable marker proteins: well annotated species (for example human and mouse vs. chicken in our study) will be easier to find markers for. We provide the markers for all the experiments available in `pRolocdata` as well as curated sets for 7 species, based on these same experiments (available in the `pRoloc` package [12]) to help researchers with their annotation. While we have shown that QSep is robust to different marker sets (figure 10), the availability of markers is a pre-requisite, and automated (i.e. that do not rely on visualising individual datasets such as documented in figure 2) and unsupervised (i.e. that do not rely on any annotation) methods, would be welcome.

**Figure 12:**
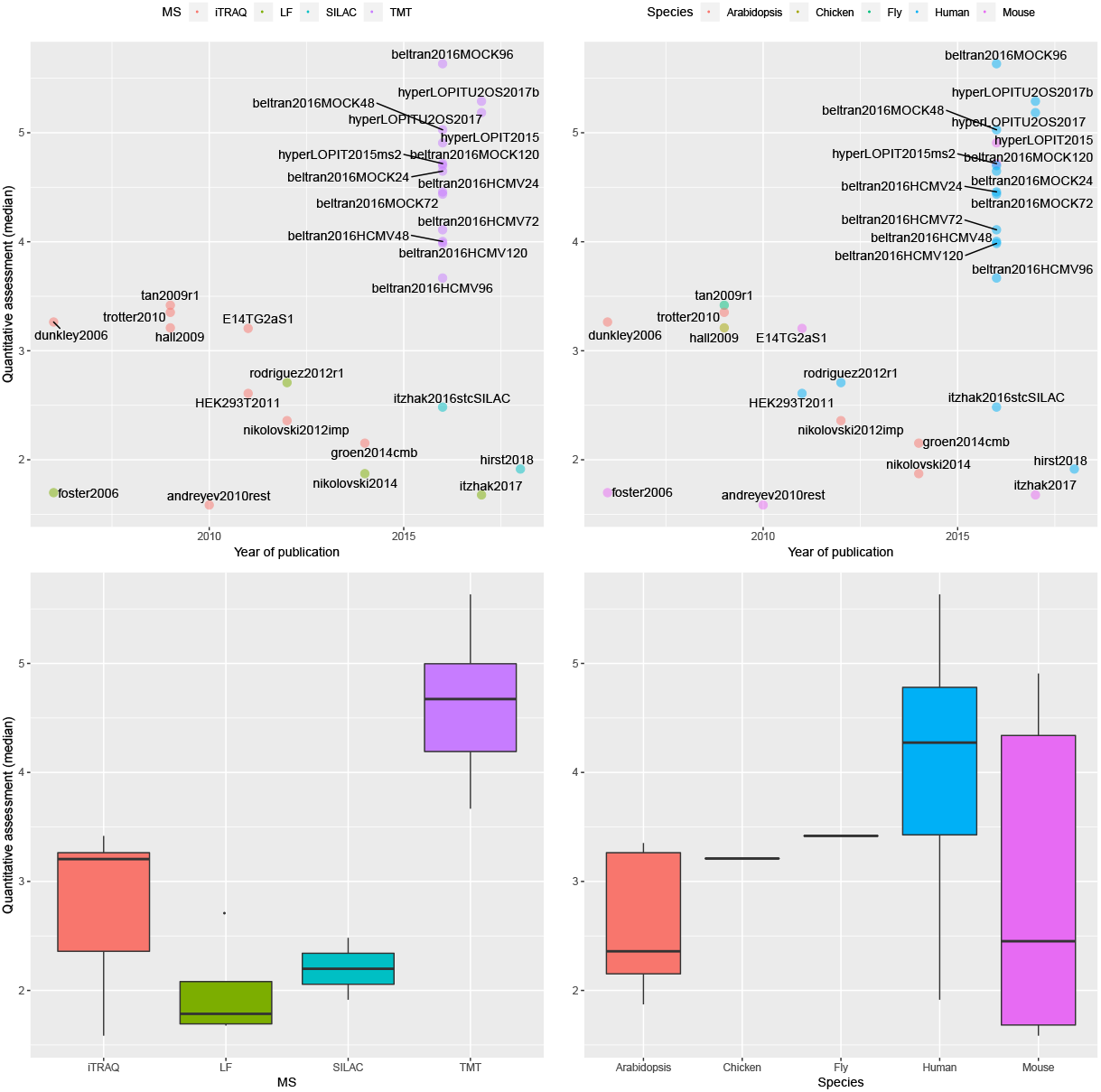
Resolution of spatial proteomics experiments over time (top panels), as presented on figure 11. Experiments are colour-coded based on the quantitation methodology (left) and species (right). The bottom panels show the median QSep resolutions for the quantitation technology (left) and species (right) respectively.

We anticipate that many laboratory-specific best practices, including various data processing and analysis steps, will influence the overall quality of the MS data and, ultimately, the resolution of the spatial proteomics experiment. As long as some best practice principles are applied (such as, for spatial proteomics in particular, adequate classification hyper-parameter optimisation and data normalisation [11]), the exact software shouldn’t matter. We however recommend the use of pRoloc [12] for robust and reproducible spatial proteomics data analysis, as it is thoroughly tested and documented, and offers a growing set of state-of-the-art visualisation and machine learning functions.

As already suggested by Lund-Johansen et al. [20], there is arguably a need for standardisation, or for general guidelines in assessing spatial proteomics data in the community? As interest in the spatial organisation of the proteome is increasing, it is imperative for the community to better define and assess the quality of spatial proteomics experiments and the reliability of protein sub-cellular assignments? The latter can assessed using improved probabilistic classifiers such as the Bayesian mixture modelling approach proposed by Crook et al. [6]. In this work, we propose the QSep metric to assess the former. Future improvements in the quality of spatial proteomics data and more reliable interpretation will be of direct benefit to the spatial proteomics researchers themselves, and will increase the usefulness of the data to the cell biology community.

## Acknowledgements

This work was supported by a BBSRC Strategic Longer and Larger grant (Award BB/L002817/1), a Wellcome Trust Technology Development Grant (Grant number 108441/Z/15/Z) and a BBSRC Tools and resources development grant (Award BB/N023129/1). The authors would like to thank Dr Claire M. Mulvey for helpful comments on the quantitative assessment.

## Author contributions

LG conceptualised the method and wrote the initial manuscript draft. LG and LMB developed the QSep code. KSL contributed datasets and feedback. All authors contributed to the manuscript and approved it.

## Appendices

## A Additional figures

This section shows the density and annotated PCA and QSep plots for the 29 datasets showcased in this manuscript.

**Figure 13:**
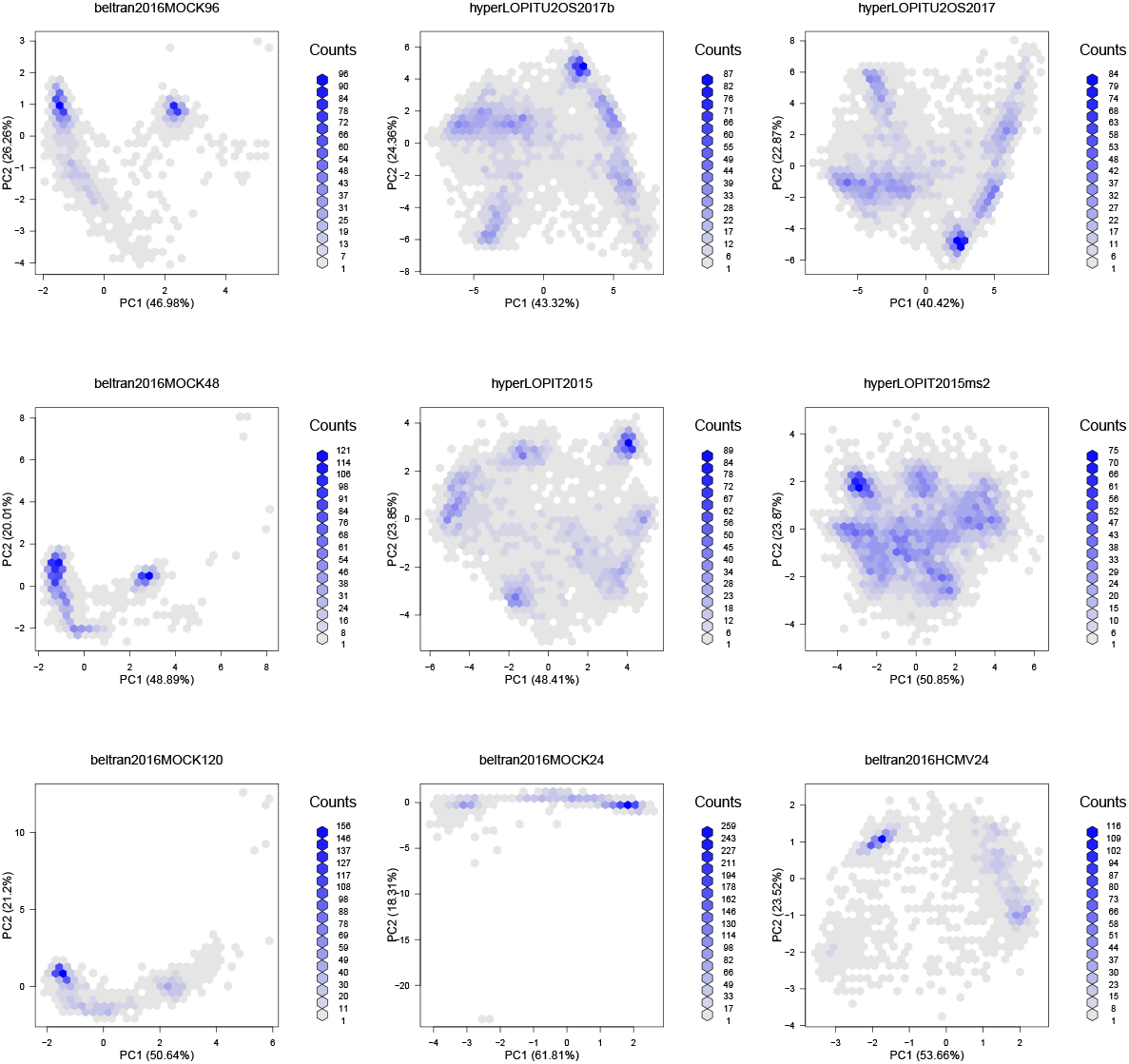
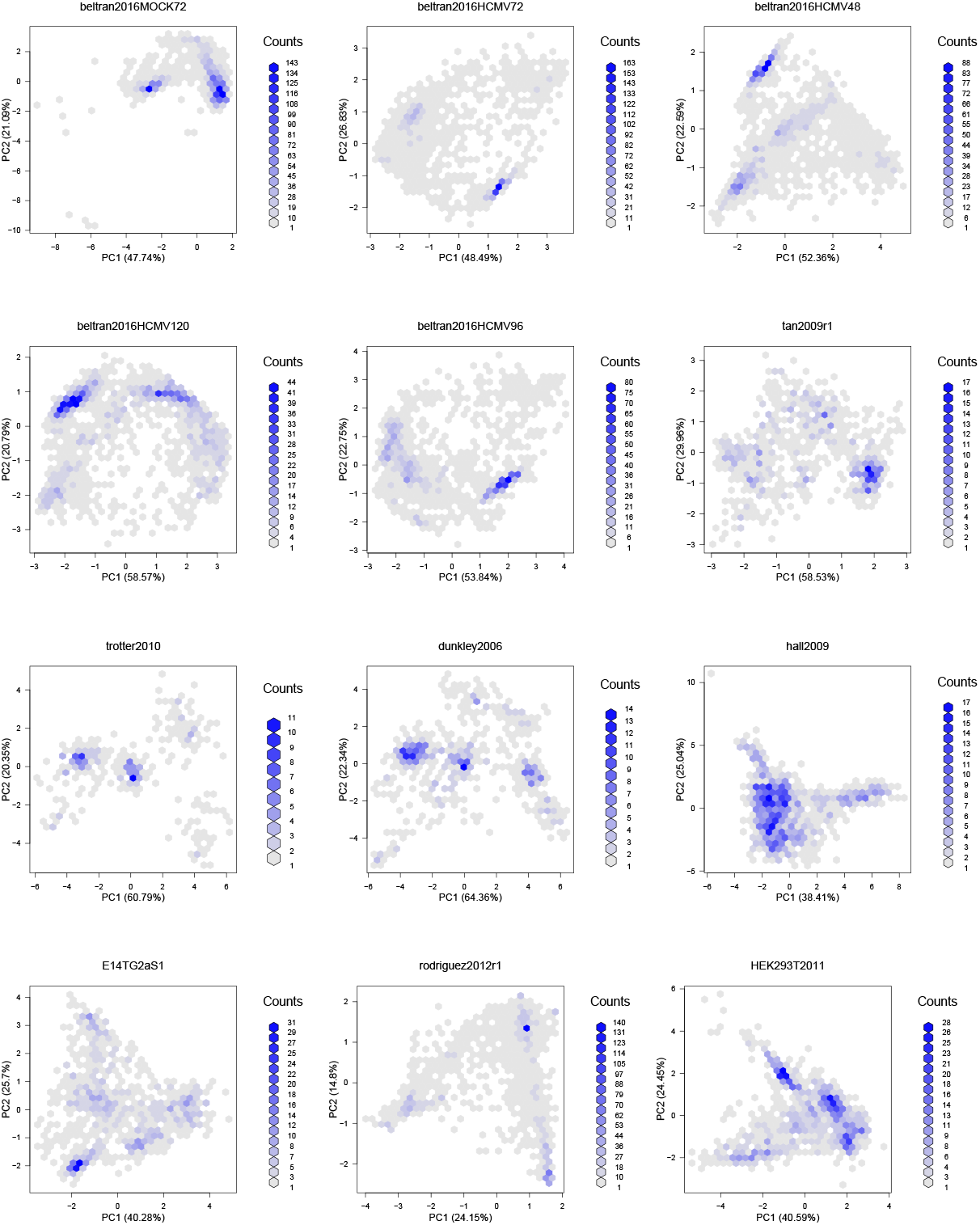
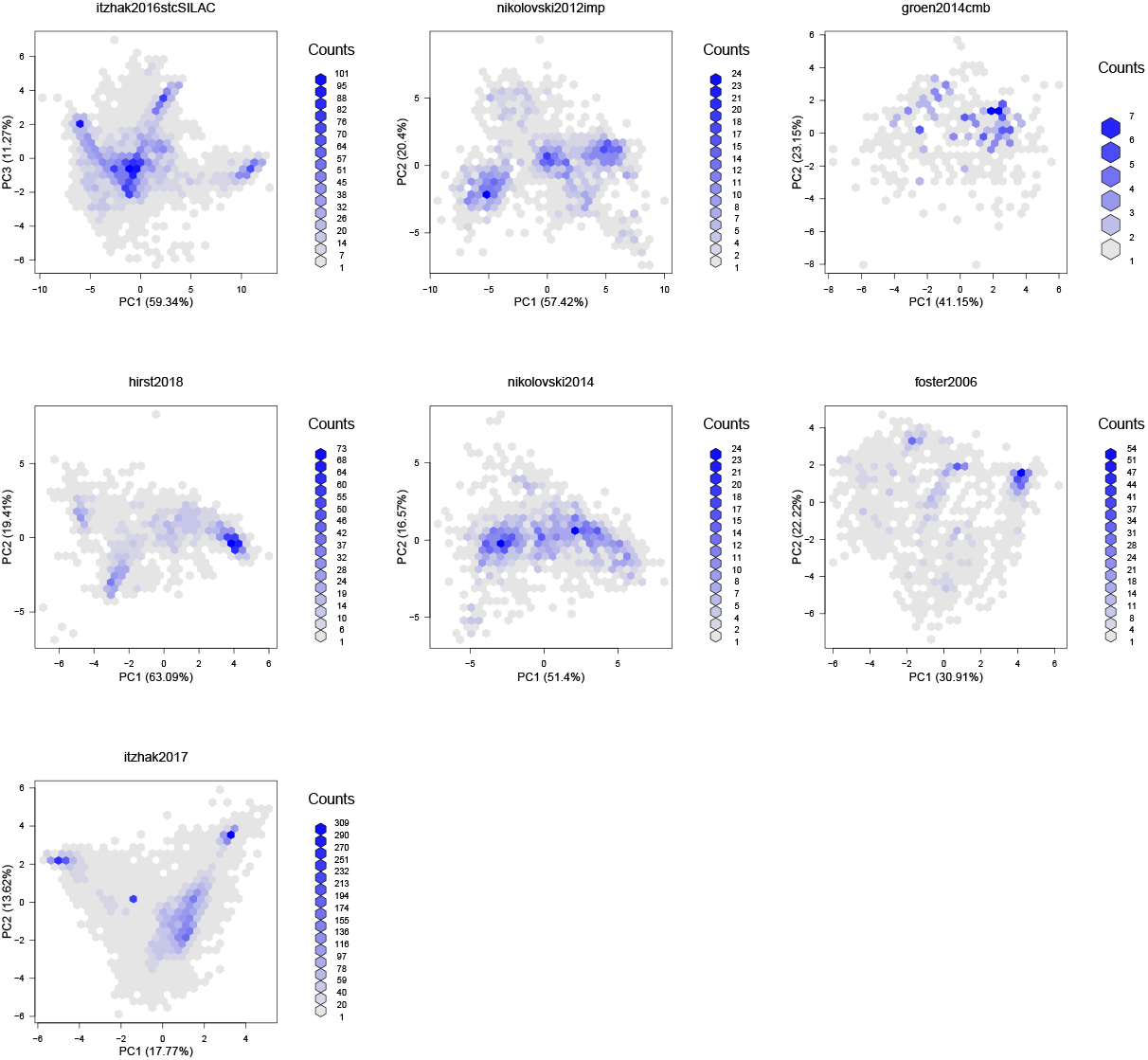
Density PCA plots for the 29 experiments used in this study. PC 1 and 2 were used except for *itzhak2016stcSILAC*, where PC 1 and 3 were used to conform to the original authors figures. The experiments are ordered according to the median average between cluster distance (see figure 8). Figures have been generated using the `plot2D` function from the `pRoloc` package.

**Figure 14:**
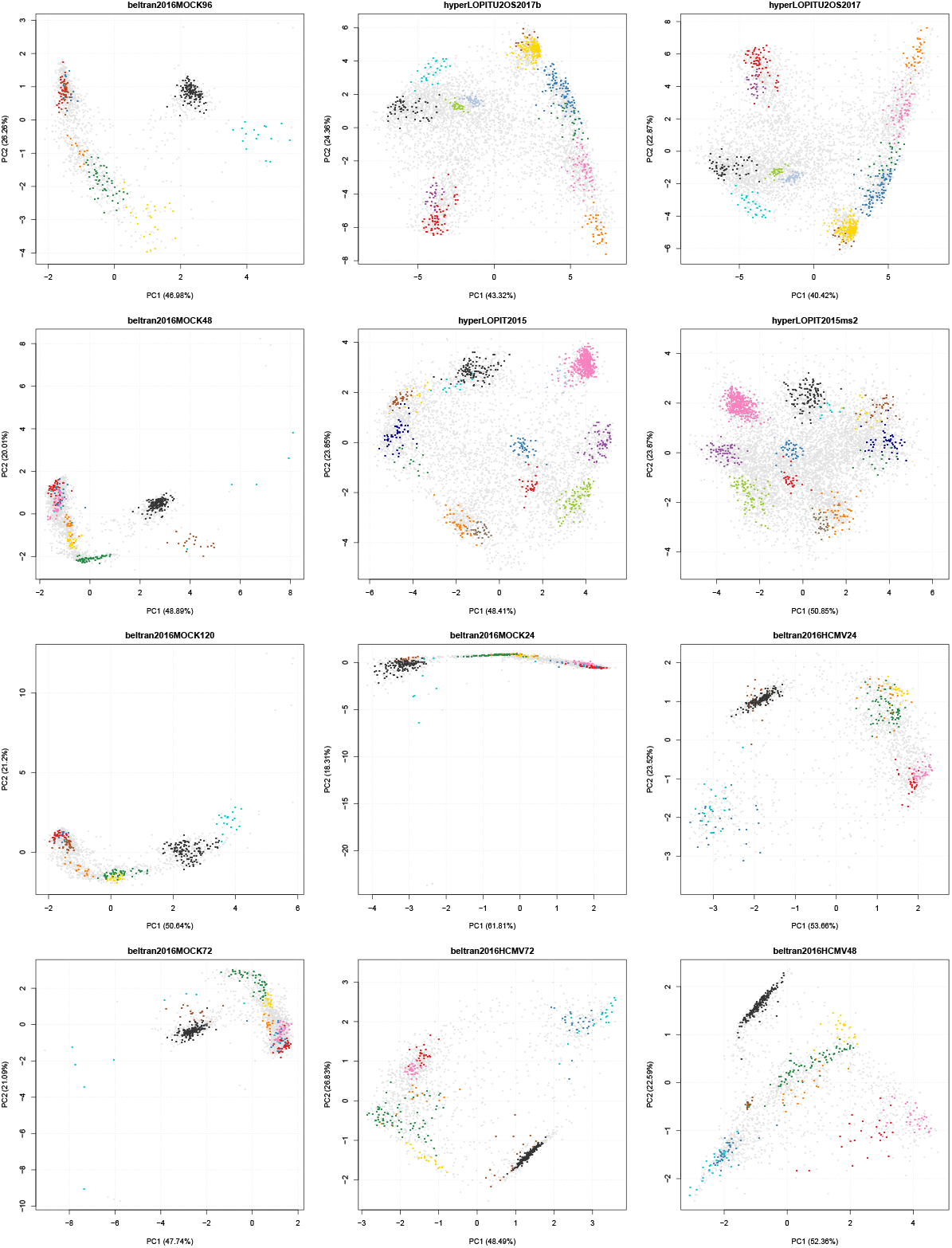
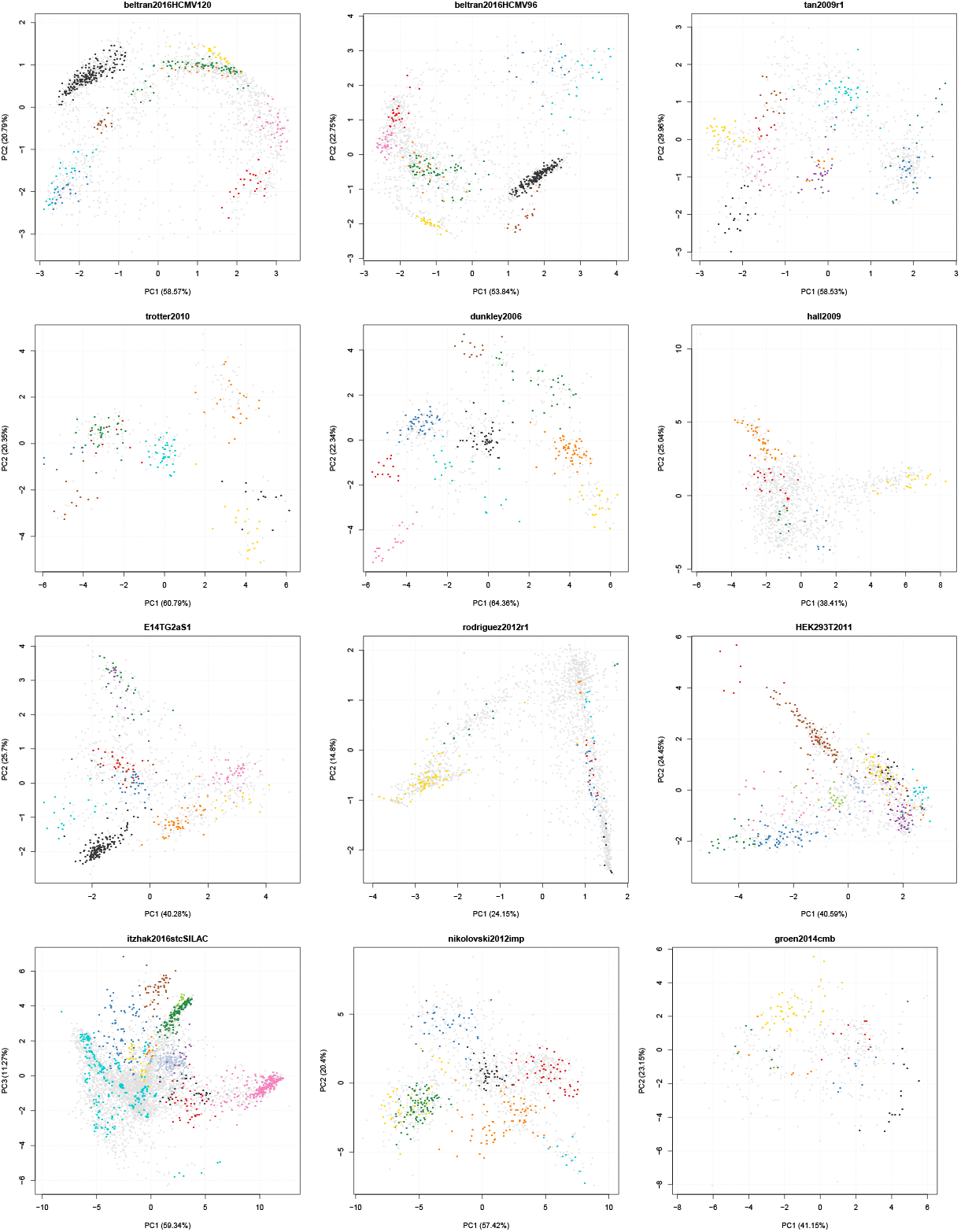
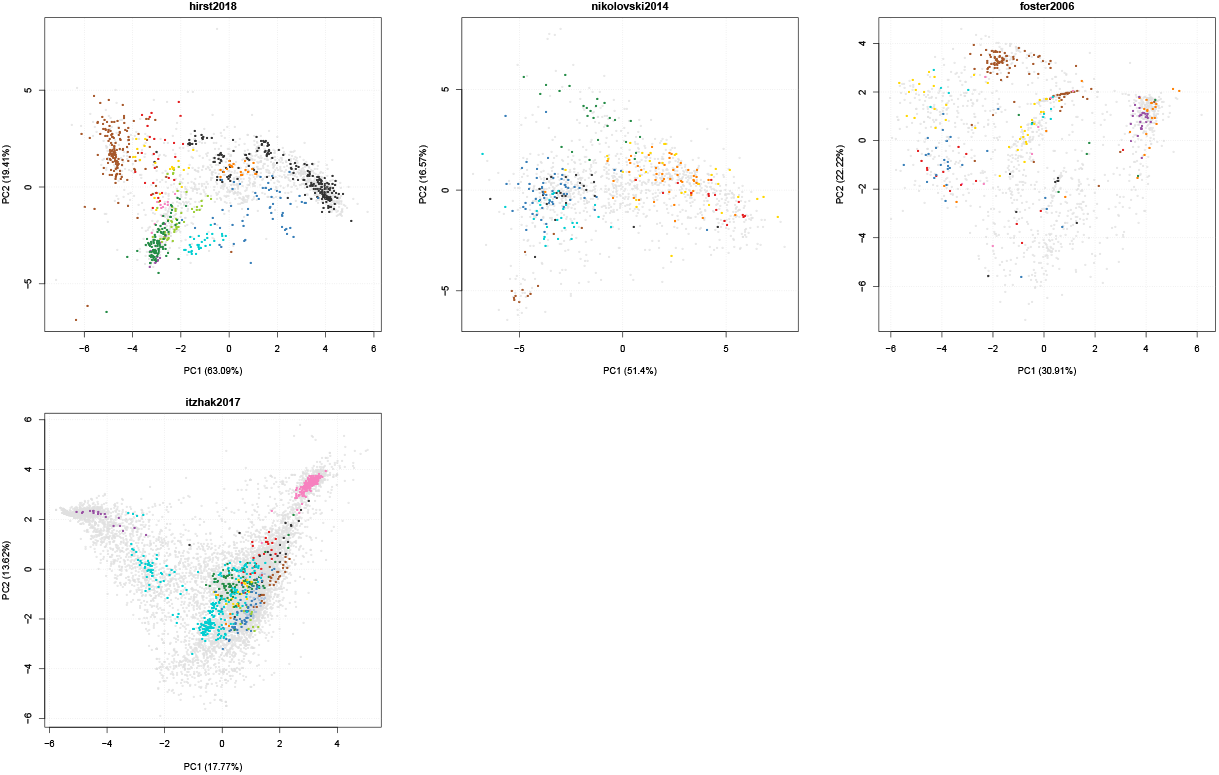
PCA plots for the 29 experiments used in this study. PC 1 and 2 were used except for *itzhak2016stcSILAC*, where PC 1 and 3 were used to conform to the original authors figures. The experiments are ordered according to the median average between cluster distance (see figure 8). The percentage of variance explained along the 2 PCs on the plots can be found in table 1. Figures have been generated using the `plot2D` function from the `pRoloc` package.

**Figure 15:**
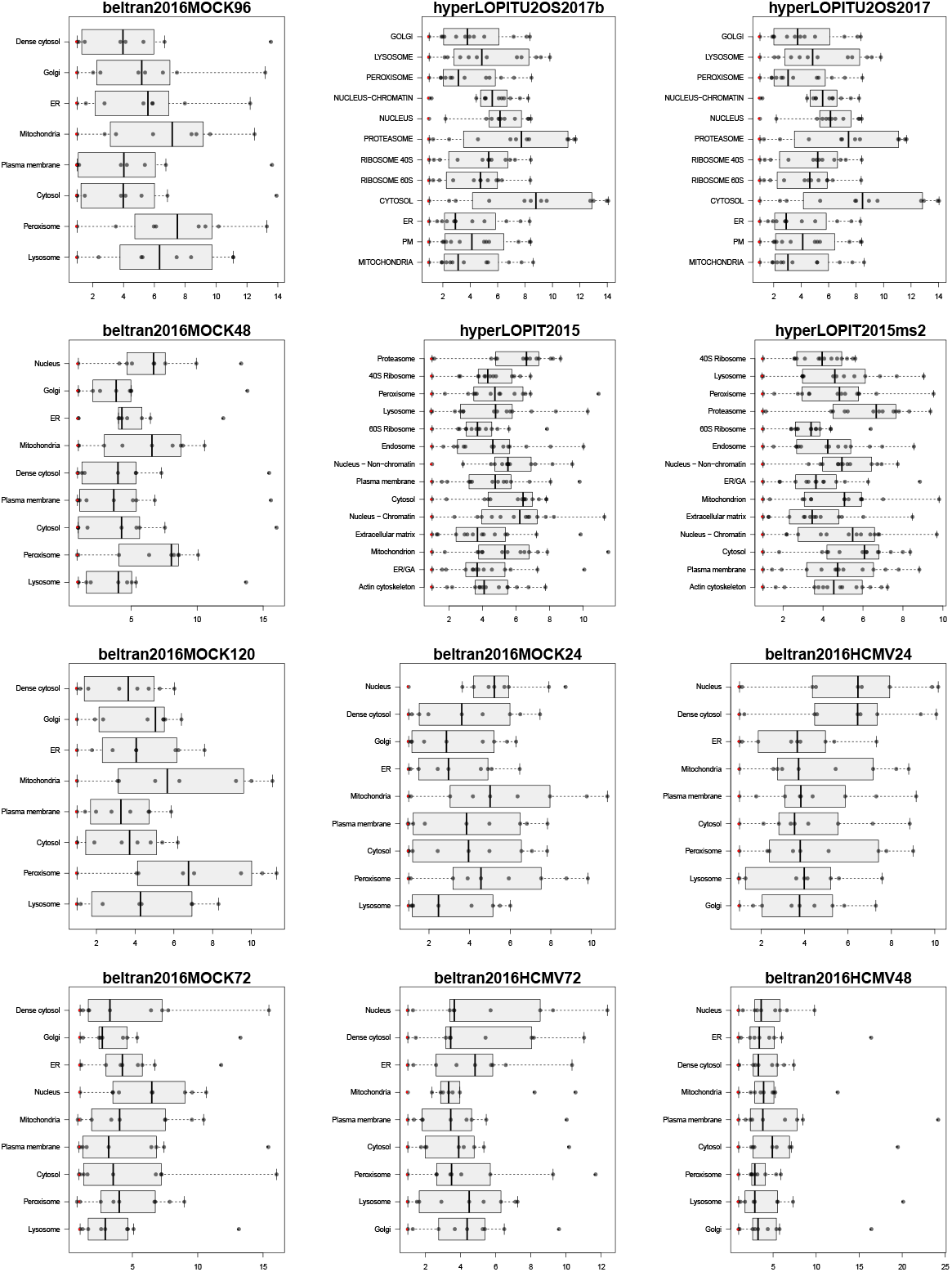
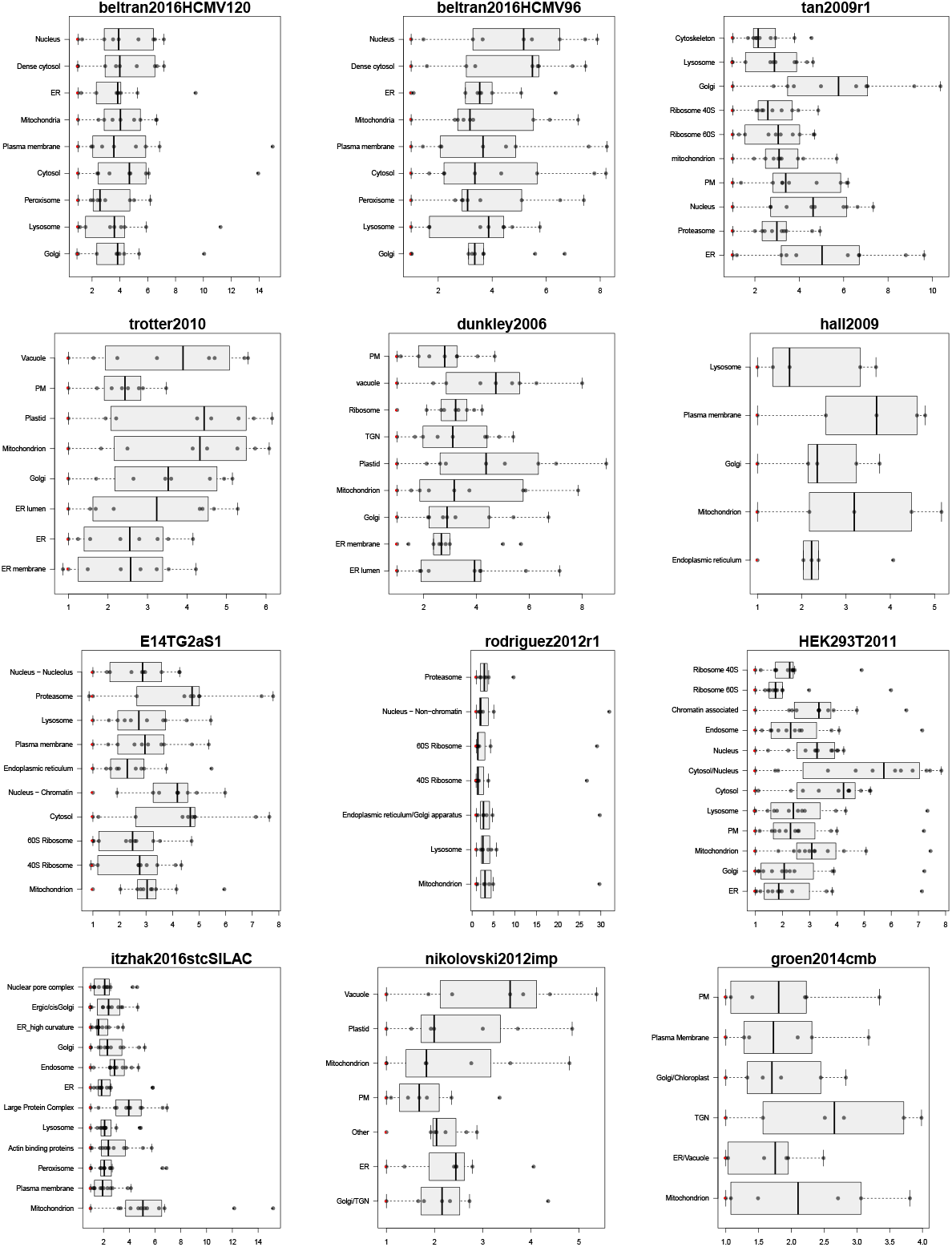
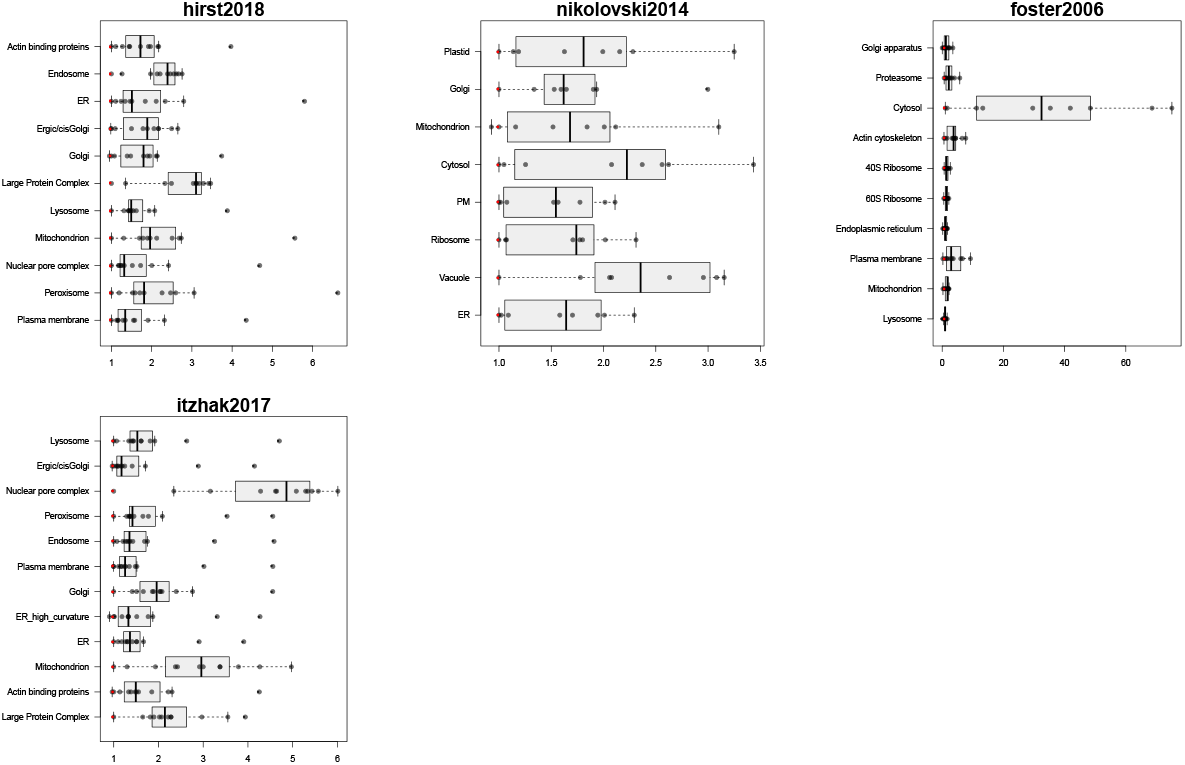
Quantitative separation boxplot for the 29 experiments used in this study. The experiments are ordered according to the median average between cluster distance (see figure 8).

**Figure 16:**
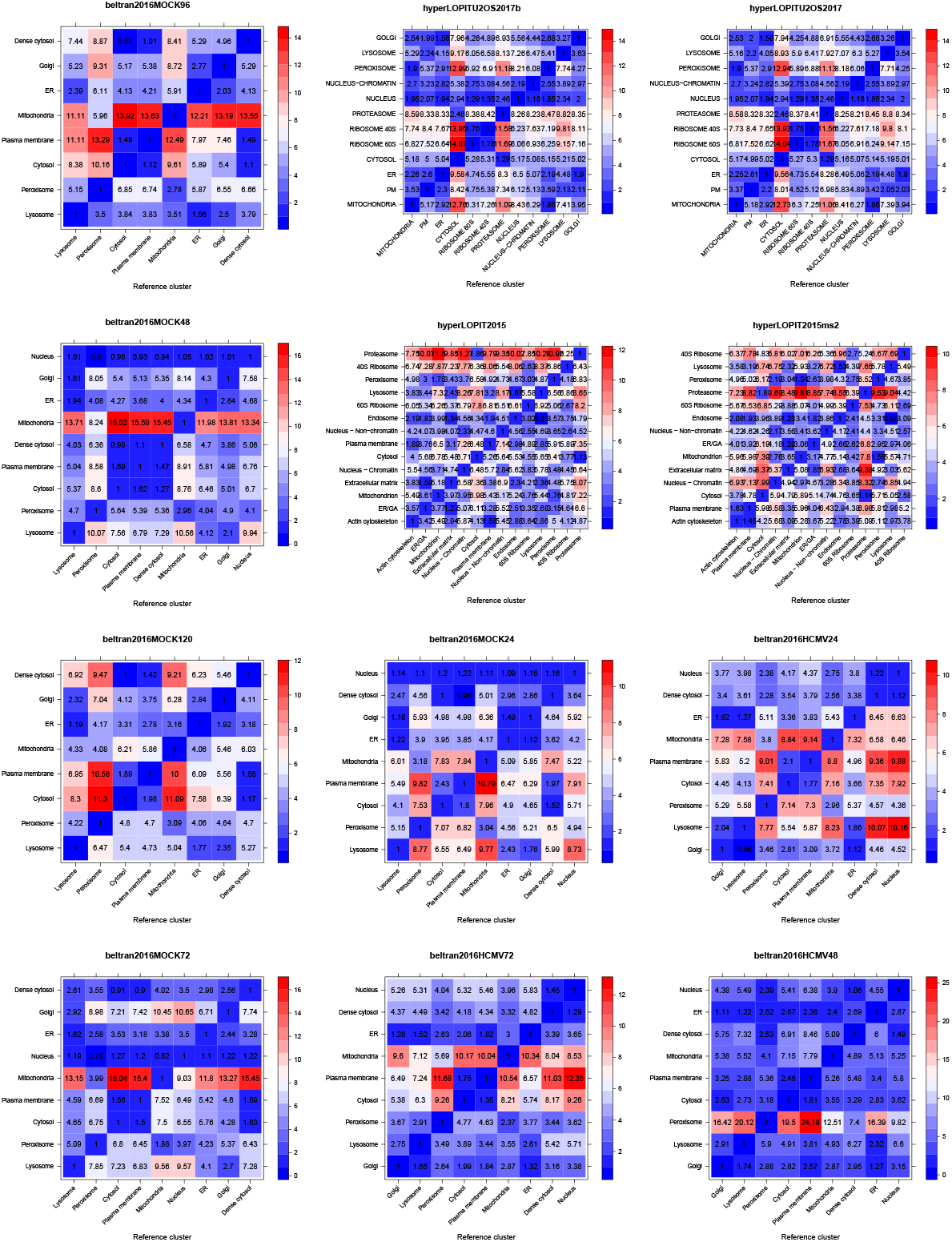
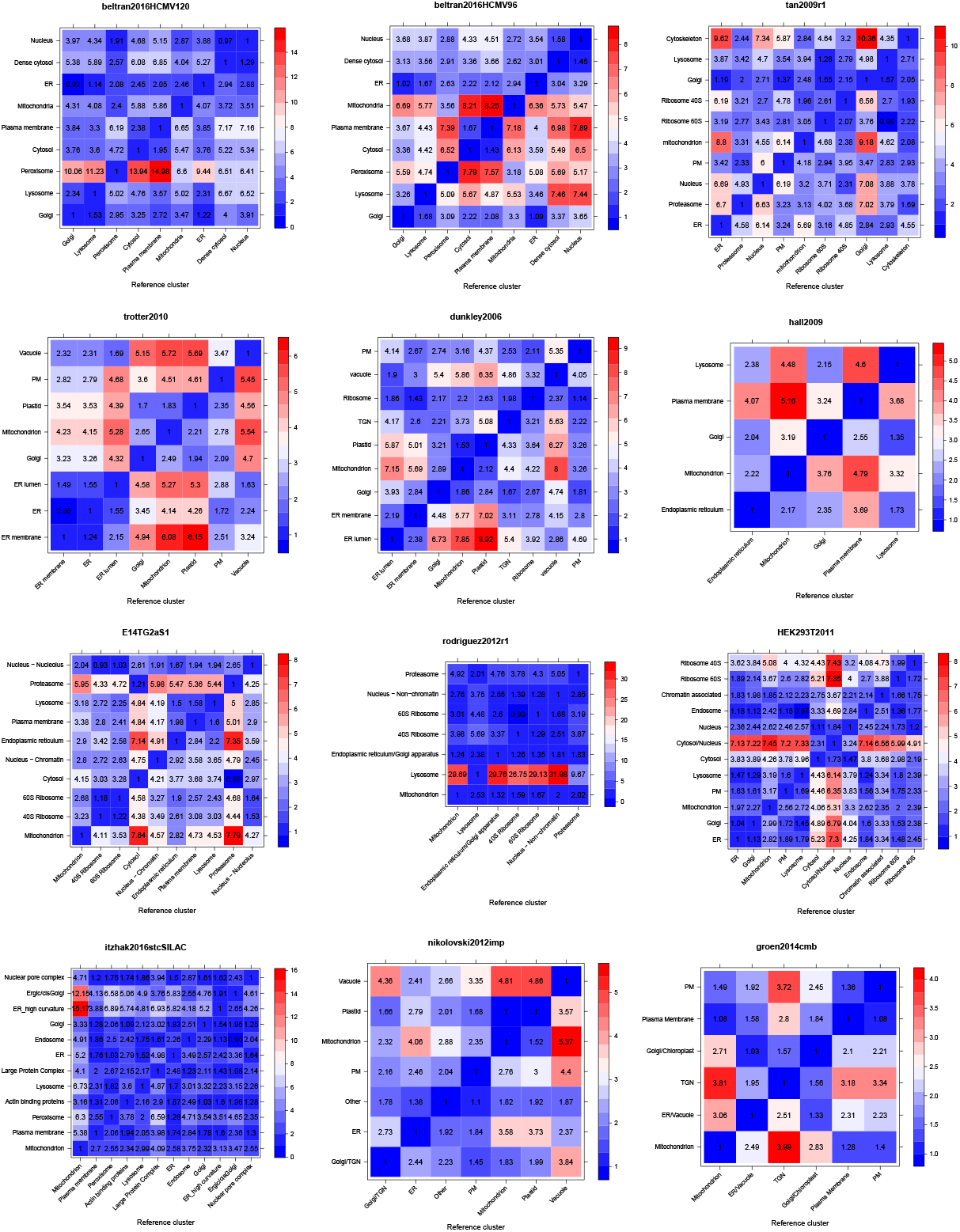
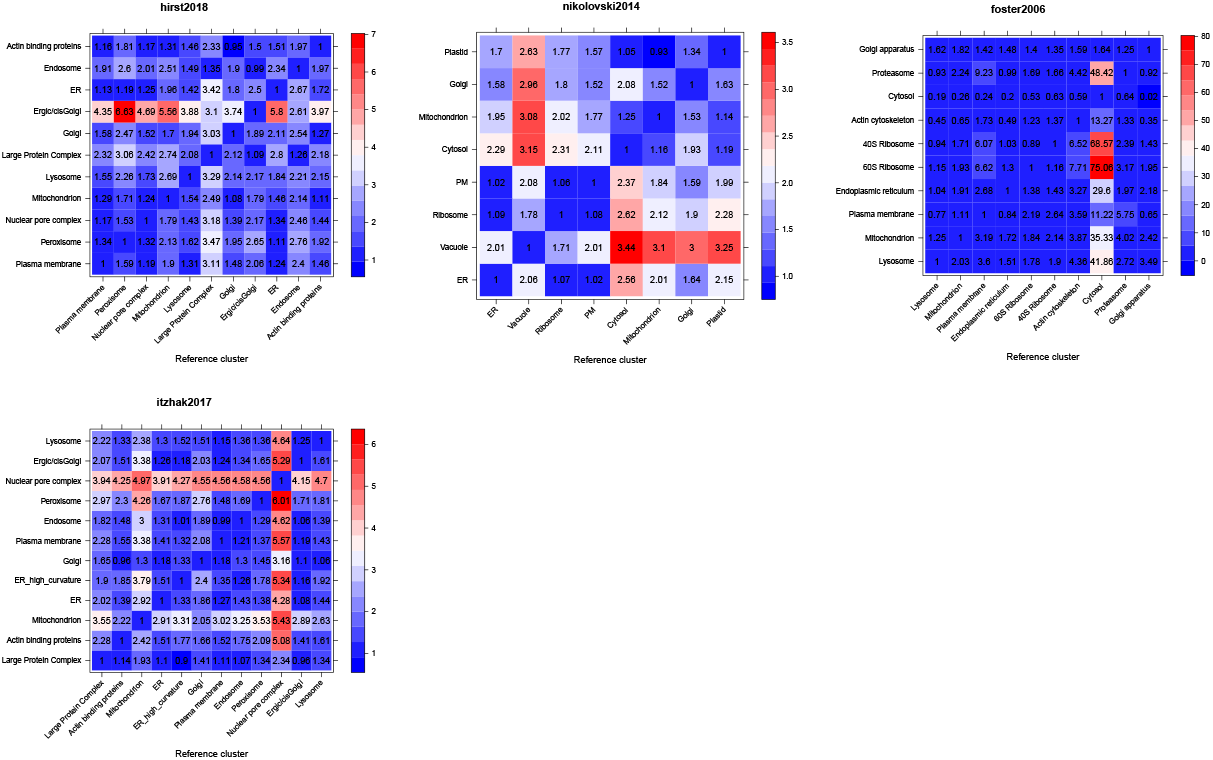
Quantitative separation heatmaps for the 29 experiments used in this study. The experiments are ordered according to the median average between cluster distance (see figure 8).

## B Session information

The software and versions used to produce this document are summarised below. The source of this document enabling to reproduce all results and figures is available in the source of this document in the public manuscript repository [9] available at https://github.com/lgatto/QSep-manuscript/.

- R version 3.5.1 Patched (2018-11-02 r75540), `x86_64-pc-linux-gnu`
- Running under: `Manjaro Linux`
- Matrix products: default
- BLAS: `/usr/lib/libblas.so.3.8.0`
- LAPACK: `/usr/lib/liblapack.so.3.8.0`
- Base packages: base, datasets, graphics, grDevices, methods, parallel, stats, stats4, utils
- Other packages: annotate 1.60.0, AnnotationDbi 1.44.0, bigmemory 4.5.33, Biobase 2.42.0, BiocGenerics 0.28.0, BiocParallel 1.16.0, cluster 2.0.7-1, ggplot2 3.1.0, ggrepel 0.8.0, hexbin 1.27.2, IRanges 2.16.0, lattice 0.20-35, MLInterfaces 1.62.0, MSnbase 2.9.2, mzR 2.16.0, patchwork 0.0.1, pRoloc 1.23.1, pRolocdata 1.20.0, ProtGenerics 1.14.0, Rcpp 1.0.0, S4Vectors 0.20.0, XML 3.98-1.16, xtable 1.8-3
- Loaded via a namespace (and not attached): abind 1.4-5, affy 1.60.0, affyio 1.52.0, assertthat 0.2.0, backports 1.1.2, base64enc 0.1-3, bibtex 0.4.2, bigmemory.sri 0.1.3, bindr 0.1.1, bindrcpp 0.2.2, BiocManager 1.30.3, biomaRt 2.38.0, bit 1.1-14, bit64 0.9-7, bitops 1.0-6, blob 1.1.1, broom 0.5.0, caret 6.0-80, class 7.3-14, coda 0.19-2, codetools 0.2-15, colorspace 1.3-2, compiler 3.5.1, crayon 1.3.4, crosstalk 1.0.0, CVST 0.2-2, data.table 1.11.8, DBI 1.0.0, ddalpha 1.3.4, dendextend 1.9.0, DEoptimR 1.0-8, digest 0.6.18, dimRed 0.2.1, diptest 0.75-7, doParallel 1.0.14, dplyr 0.7.7, DRR 0.0.3, e1071 1.7-0, evaluate 0.12, flexmix 2.3-14, FNN 1.1.2.1, foreach 1.4.4, fpc 2.1-11.1, gbm 2.1.4, gdata 2.18.0, genefilter 1.64.0, geometry 0.3-6, ggvis 0.4.4, glue 1.3.0, gower 0.1.2, grid 3.5.1, gridBase 0.4-7, gridExtra 2.3, gtable 0.2.0, gtools 3.8.1, highr 0.7, hms 0.4.2, htmltools 0.3.6, htmlwidgets 1.3, httpuv 1.4.5, httr 1.3.1, hwriter 1.3.2, igraph 1.2.2, impute 1.56.0, ipred 0.9-8, iterators 1.0.10, kernlab 0.9-27, knitr 1.20, labeling 0.3, LaplacesDemon 16.1.1, later 0.7.5, lava 1.6.3, lazyeval 0.2.1, limma 3.38.2, lpSolve 5.6.13, lubridate 1.7.4, magic 1.5-9, magrittr 1.5, MALDIquant 1.18, MASS 7.3-51.1, Matrix 1.2-14, mclust 5.4.1, memoise 1.1.0, mime 0.6, mixtools 1.1.0, mlbench 2.1-1, ModelMetrics 1.2.2, modeltools 0.2-22, munsell 0.5.0, mvtnorm 1.0-8, mzID 1.20.0, ncdf4 1.16, nlme 3.1-137, NMF 0.21.0, nnet 7.3-12, pcaMethods 1.74.0, pillar 1.3.0, pkgconfig 2.0.2, pkgmaker 0.27, pls 2.7-0, plyr 1.8.4, prabclus 2.2-6, preprocessCore 1.44.0, prettyunits 1.0.2, prodlim 2018.04.18, progress 1.2.0, promises 1.0.1, proxy 0.4-22, purrr 0.2.5, R6 2.3.0, randomForest 4.6-14, RColorBrewer 1.1-2, RcppRoll 0.3.0, RCurl 1.95-4.11, rda 1.0.2-2.1, recipes 0.1.3, registry 0.5, reshape2 1.4.3, rlang 0.3.0.1, rngtools 1.3.1, robustbase 0.93-3, rpart 4.1-13, RSQLite 2.1.1, sampling 2.8, scales 1.0.0, segmented 0.5-3.0, sfsmisc 1.1-2, shiny 1.2.0, splines 3.5.1, stringi 1.2.4, stringr 1.3.1, survival 2.43-1, threejs 0.3.1, tibble 1.4.2, tidyr 0.8.2, tidyselect 0.2.5, timeDate 3043.102, tools 3.5.1, trimcluster 0.1-2.1, viridis 0.5.1, viridisLite 0.3.0, vsn 3.50.0, whisker 0.3-2, withr 2.1.2, zlibbioc 1.28.0

## Footnotes

1 This results from the fact that they used a simple distance measurement, termed *χ*^2^, against very few markers to base their sub-cellular localisation prediction

2 This number is relatively low, and we would typically recommend at least 13 markers per class to perform cross-validation when optimising classifier parameters. See Gatto et al. [11] and the main `pRoloc` tutorial for details.

